# Male contributions to female-male mutualism overcome the twofold cost of sex

**DOI:** 10.1101/2024.01.11.575303

**Authors:** Yukio Yasui, Eisuke Hasegawa

## Abstract

The increasing rate of anisogamy (fusion between eggs and sperm) against asexual thelytoky is necessarily halved under a 1:1 sex ratio because males lay no eggs. The present dominance of anisogamy despite this twofold cost of male production is the greatest enigma in evolutionary biology. Various benefits of sex are insufficient to overcome this cost. How can anisogamy be maintained? Here, we propose hypotheses that anisogamy evolved as a female-male mutualism in which females produce large eggs to ensure zygote development and males produce many sperm to ensure fertilisation. In the real world with finite resources, thelytoky cannot express its twofold increasing potential, and various male functions reduce the thelytoky’s performance. Thus, the twofold cost is an artefact of the infinite resource assumption. Simple mathematical models and simulations showed that when males and thelytokous females are equally competitive and acquire the same amount of resources but males can develop with fewer resources than females, anisogamy outcompetes thelytoky. When males need more growth costs than females, as in sexually selected species, the excess mating activity of “harmful” males reduces the fitness of thelytokous females. The significance of sexes, a question since Aristotle’s time, is explained by male contribution to sustaining mutualism.

## Main text

Sex is the exchange of parts of genomes between individuals^1,2^. Gametic sexual reproduction dominates in eukaryotes^1^. In this mode, both parents produce gametes with half of their genomes via meiosis and fuse two gametes into a zygote, restoring the full genomes. In contrast, asexual reproduction transmits all genomes to offspring without meiosis. This half rate of genome transmission in gametic sexual reproduction is referred to as the “cost of meiosis”^2,3^. In most multicellular sexual organisms, females and males produce large eggs and tiny sperm, respectively^1,2,4,5^. This gamete size dimorphism is referred to as “anisogamy”, in which resource investment in a gamete is different but the genetic contributions are equal between sexes. If the parents produce the same number of males as females (birth numerical sex ratio, female:male = 1:1), the rate of increase of the anisogamous population will be half that of the asexual rival (thelytoky) because males do not lay eggs. This reduced productivity in anisogamy is called the “cost of male production”^2,3,6,7^.

These costs are generally known as the “twofold cost of sex”^1-3,6,8^. Previous authors^3,9^ suggested that the cost of meiosis disappears from the sexual population once the sex alleles are fixed at the sex-controlling locus (loci) because meiosis cannot dilute the fixed locus. However, our recent study^10^ has shown that this fixation of the sex alleles must have occurred mechanically during the first sexual reproduction, making isogamy much easier to evolve than previously thought.We^10^ also suggested that other loci in the genome are forced to follow the sex locus to avoid extinction by the Muller’s ratchet (accumulation of deleterious genes)^9^ and ensure evolvability. In contrast, the evolution and maintenance of anisogamy is still a mystery because male production imposes *twofold costs per generation*, under which the population size of thelytoky is 2^10^ = 1024 times larger than that of anisogamy after 10 generations. It seems impossible to find some immediate benefits of sex that would outweigh this enormous difference. Various traditional hypotheses, such as the Fisher-Muller effect^11,12^, the Red Queen hypothesis^13-15^, Muller’s ratchet^9^ and the Kondrashov effect^16^, are too weak and too slow. Hence, the protection of anisogamy from thelytoky invasion is the true problem regarding the cost of sex.

Here, we present two general conditions that make anisogamy so successful: 1) the female-male relationship becomes a mutualism, and 2) males contribute to this mutualism by preventing thelytoky invasion. For 1), we show that both sexes help each other; males produce many sperm to ensure fertilisation of eggs, and females produce large, nutritious eggs to ensure zygote development. This mutualism is the main force that has led to the evolution of anisogamy from isogamous ancestors and is still maintained in all anisogamous species. For 2), we introduce a sustainability perspective^17^; under finite resources, low-productivity, low-consumption strategies will persist longer than high-productivity, high-consumption strategies (see below). Based on this perspective, we propose two hypotheses regarding male contributions. 2a) The “light eater male” hypothesis, in which males save their own resource consumption and the unused resources are used for female reproduction. When this condition is met, thelytoky individuals deplete their resources and go extinct earlier than anisogamy. 2b) The “harmful male” hypothesis, in which highly sexually active males, evolved through sexual selection and sexual conflict, forcibly copulate with thelytokous females, reducing the thelytoky fitness^18-20^.

### General condition 1: Anisogamy as a mutualism

Isogamy, as the ancestral form of gametic sexual reproduction, is a mutualism because both parents invest equal amounts of nutrition into the zygotes^3,10^. Traditional twofold cost theories^21,22^ treat derived anisogamy as male parasitism on female reproductive investment simply because a single sperm is much smaller than an egg. However, we consider that anisogamy is still mutualism, but females and males invest in different ways. Sperm are produced in large numbers instead of being small and used in large quantities at once^23^. Only one of these sperm results in an embryo, and the rest die and are wasted. This large amount of wasted sperm should be considered a hidden investment by the male. In other words, males invest in the prezygotic process (ensuring fertilisation), and females invest in the postzygotic process (ensuring survival). Therefore, the traditional parasitic view^22^ is incorrect because it overlooks prezygotic investment by males. In truth, anisogamy is a mutualism from the beginning because females and males have been compensating for each other’s weaknesses.

### Significance of males: Many sperm resolve the gamete limitation problem

Complex multicellular organisms require many resources (large zygotes) for ontogeny^10^, but large, immobile isogametes are unlikely to encounter each other^5^. In addition, large amounts of cytoplasm prevent fusion (imagine that whether two half-sized ostrich eggs can fuse with each other). Thus, anisogamy had to evolve^10^. The weakness for females is that large, immobile eggs tend to die without being fertilised. To solve this “gamete limitation problem”^5,6^, males disperse large numbers of sperm^5^ and ensure egg fertilisation, assuming that most of them will be wasted. The weakness for males is that small sperm cannot nourish the zygote, even if it successfully fertilises eggs. To solve this problem, females invest nutrition in large eggs in advance. A positive feedback escalates between more sperm and larger eggs, and the transition from isogamy to anisogamy occurs very quickly^5,6,10,24,25^.

### Invasion of costly anisogamy into an isogamous population

In the early stages of anisogamy evolution, sperm production (number of spermatocytes × 4) was still low, so males could fertilise the eggs of only a few females. However, as multicellularisation progressed^10^, males increased the number of spermatocytes and could potentially fertilise the eggs of many females. Males compete with each other for mating opportunities, and the winner achieves multiple matings (polygamy)^21^. This male reproductive success feeds back to his mother (Fisher’s son’s effect^11,21^), maintaining mutualism.

Our simulation (Fig. 1) confirmed this scenario. When a mutant allele Y (male allele) that produces sperm enters a population of allele X that produces isogametes, the frequency of Y increases rapidly owing to the fertilisation advantage of sperm over isogametes and the mechanical necessity that the XX females that mate with XY males produce XY sons and XX daughters at a 1:1 ratio. These sperm-derived females thereafter perform anisogamy and bear the twofold cost of male production, but this does not stop the rapid spread of the Y allele because the frequency increase in Y in a gene pool is faster than the expansion of the gene pool itself^22^. In other words, it is more advantageous for the X allele to be fertilised by Y sperm and become an XY male than to fuse with the isogamete X and remain an isogamous female. The X allele hitchhikes the Y allele’s ability to spread, and the mother increases the frequency of her genes through her son (Fig. 1c).

**Fig. 1.**
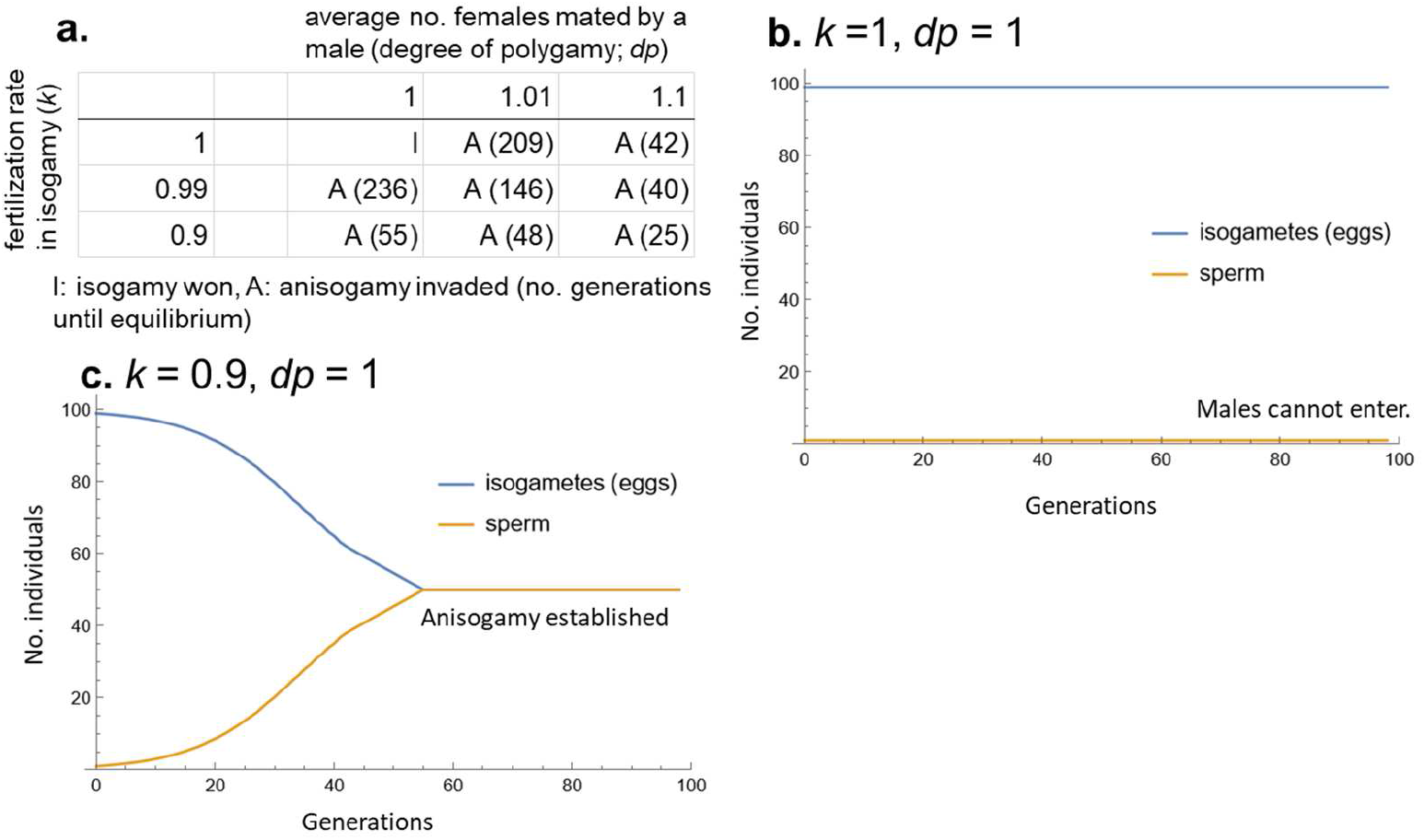
Results of simulation 1. Invasion of the Y allele (male) into an isogamous population (genotype XX) (population size 100). In the early generations, an invading male has great mating success, but this success gradually decreases as the number of males increases over generations. **a**. Although isogamy produces twice as many daughters as anisogamy (4 in isogamy and 2 in anisogamy), isogamy is quickly replaced by anisogamy if a single male can fertilise isogametes from, on average, slightly more than one female or if the sperm slightly improves the fertilisation rate compared to isogamy. **b**. If there is no fertilisation failure in isogamy and males mate with only one female, invasion of the anisogamous allele is impossible. **c**. If sperm improve the fertilisation rate by 11% (= 1/0.9) compared to isogamy, anisogamy evolves even in monogamy. *dp*: average number of mates per male (degree of polygamy). *k*: relative fertilisation rate in isogamy (*k* = 1 in anisogamy).

## General condition 2: Male contribution

### a) The “light eater male” hypothesis

To achieve a 1024-fold increase, thelytoky requires a 1024-fold amount of resources. Where will it be supplied from? The traditional twofold-cost theory implicitly assumes infinite resources, but in fact, resources are finite in nature. Under finite resources, thelytokous females do not always achieve their twofold intrinsic rate of increase. Instead, the low increasing rate due to male production may favour anisogamy. In the following sections, we examined this possibility in several conditions.

#### Model 1: Scramble competition between offspring clutches

Anisogamous females, males and thelytokous females (derived species) are assumed. The offspring compete for the resources needed for their own development. Imagine moth larvae laid in egg masses on host plants. These larvae feed in sibling groups and, as they grow, meet other family groups (including anisogamy and thelytoky) and compete for food. Competition is of the scramble type, in which competitors do not fight each other but compete to exploit finite resources more quickly. Three types of individuals of the same age are equally competitive. Every generation is supplied with *R* units of resources, meaning the environmental carrying capacity is *R* individuals. In the initial generation, the numbers of adult sexual females, males and thelytokous females are *f*_0_, *m*_0_ and *t*_0_, respectively. Because of the 1:1 sex ratio, *m*_0_ = *f*_0_. When a sexual female produces *n* daughters + *n* sons and a thelytokous female produces 2*n* daughters (i.e., the twofold cost in anisogamy), the total resource of the population is allocated to each larval individual by 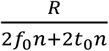. The amounts of resources required for the growth of one juvenile sexual female, one male and one thelytokous female are *d*_*f*_, *d*_*m*_ and *d*_*t*_, respectively. Considering how many pairs of females and males can be grown with the resources allocated to the two individuals under a sex ratio of 1:1, the potential rate of increase (= fitness) in the sexual population is

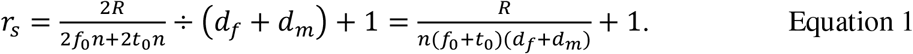

Considering how many thelytokous females can be grown with the resources allocated to one juvenile, the potential rate of increase of the thelytokous population is

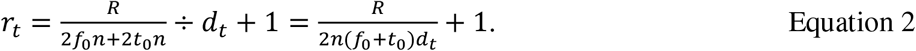

+1 is the measure to keep the increasing tendency of each type (without this value, the population does not reach carrying capacity). The condition under which the rate of increase in sexuals exceeds that of thelytoky is

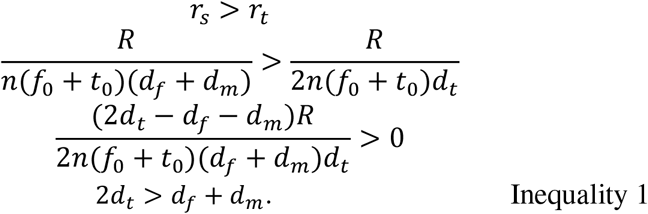

Inequality 1 is satisfied when *d*_*t*_ > *d*_*m*_ > 0 because we assume *d*_*f*_ = *d*_*t*_. This implies that if males consume fewer resources (growth costs) than sexual and thelytokous females, anisogamy wins thelytoky. For example, if *d*_*f*_ = *d*_*t*_ =1, *d*_*m*_ = 0.9 and the population reached carrying capacity (*R* = 100, *f*_0_ = 25, *t*_0_ = 50, *n* = 2), the increasing rates are *r*_*s*_ = 1.351 and *r*_*t*_ = 1.333. On average, 33.77 sexual pairs and 66.67 thelytokous females (134 in total) were born, and *f*_1_ = 25.16 pairs and *t*_1_ =49.67 (100 in total), respectively, survived after equal competition. Consequently, only anisogamy could increase its frequency because resources secured but not eaten by males increased the survival of other sexual juveniles. In the moth case where a clutch of larvae is equally competitive between anisogamous and (conjectural) thelytokous species, anisogamous larval clutches usually contain both sexes, so food left uneaten by males will be eaten by their sisters immediately adjacent to them rather than the thelytokous larvae in a separate clutch. Simulations (Fig. 2a) show that thelytoky increased much faster than anisogamy at low densities, but once the population reached the carrying capacity, thelytoky stopped increasing, and anisogamy gradually increased in frequency, eventually occupying the population.

**Fig. 2.**
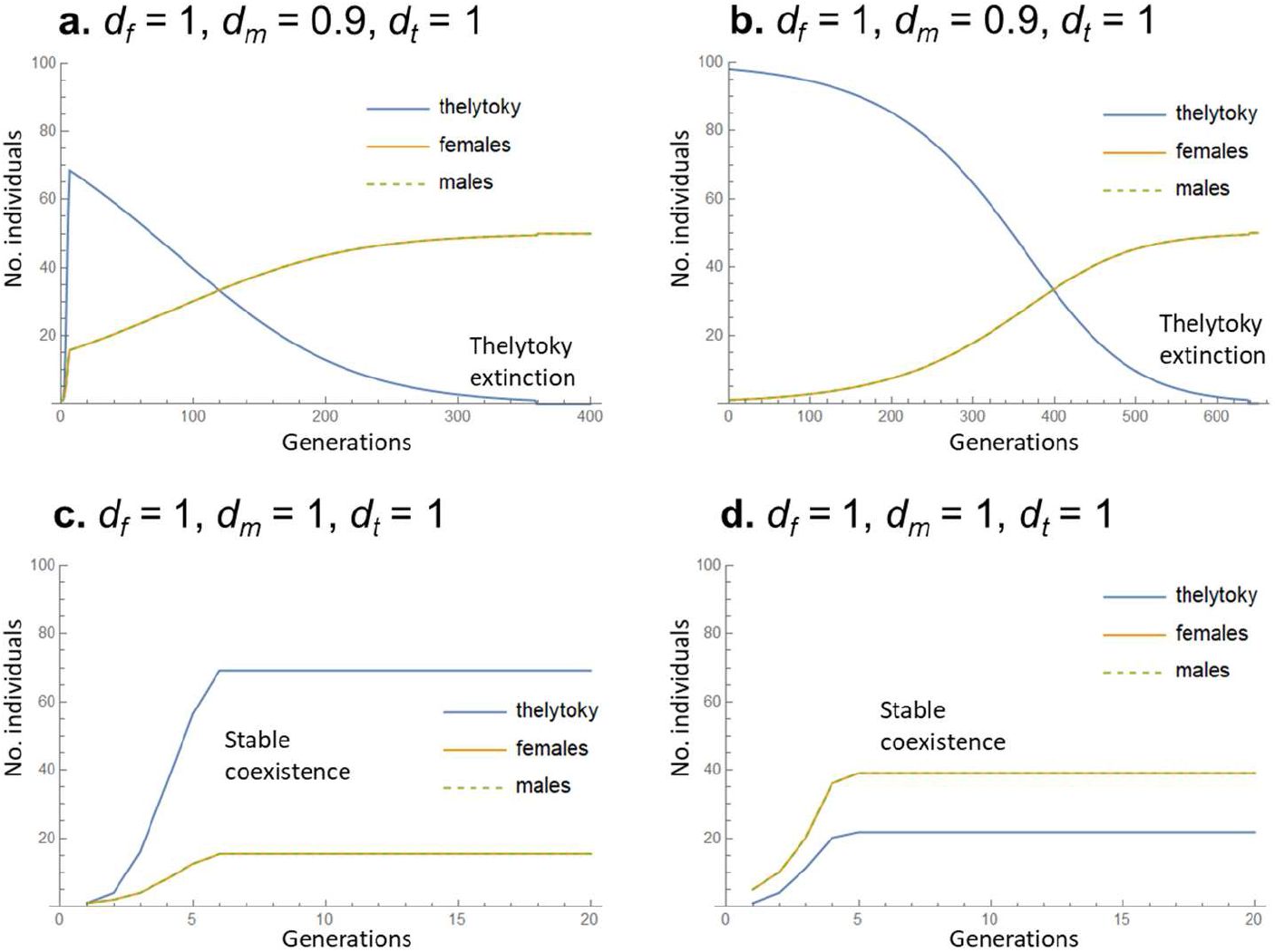
Results of simulation 2. Less-consuming males win thelytoky. Resource competition under Model 1 settings was simulated (see Model 1). No. offspring produced per mother at low density: 2 females + 2 males in anisogamy, 4 females in thelytoky. This determines the upper limit of the increasing rate: If *r*_*s*_ > 2, then *r*_*s*_ = 2, and if *r*_*t*_ > 4, then *r*_*t*_ = 4. **a-b**. Growth cost is lower in males than females (*d*_*f*_ = *d*_*t*_ = 1 and *d*_*m*_ = 0.9). **a**. When an anisogamous pair and a thelytokous female entered an empty environment (*f*_0_ = 1 and *t*_0_ = 1), thelytoky initially increased rapidly, but when the population reached carrying capacity (*R* = 100), thelytoky began to decline and became extinct, while anisogamy continued to increase and reached fixation. **b**. Anisogamy was able to invade the population occupied by the thelytoky (*f*_0_ = 1, *t*_0_ = 98). **c-d**. When the growth cost was equal between sexes (*d*_*f*_ = *d*_*t*_ = *d*_*m*_ = 1, meaning equal increasing rate *r*_*s*_ = *r*_*t*_), initial conditions (*f*_0_ = 1, *t*_0_ =1 in **c** and *f*_0_ = 5, *t*_0_ = 1 in **d**) determined the composition of the community at saturation, but once saturation was reached, it was maintained because of the settings where drift did not act. Lines of anisogamous females and males overlap because of the 1:1 sex ratio. See text.

#### Model 2: Adults compete for resources, within which their offspring grow up

The numbers of adult individuals of anisogamous females, males and thelytokous females are *f, m* and *t*, respectively. We assume that males are lighter in body weight but more aggressive than females (body length does not differ). As a result, adults are equally competitive in contest competition, so the resource is divided between the three parties into 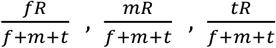 respectively. Sexual females and males form monogamous pairs and reproduce on the basis of resources 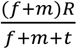 acquired by the pair. To grow to adulthood, one juvenile sexual female, one male and one thelytokous female require resources *d*_*f*_, *d*_*m*_ and *d*_*t*_, respectively. The number of offspring reaching adulthood in a sexual population is 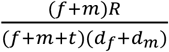 pairs at a sex ratio of 1:1.The number of offspring reaching adulthood in the thelytokous population is 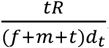.The rate of increase per generation (= fitness) is

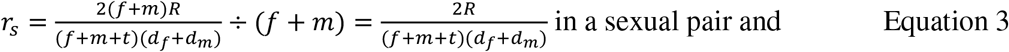

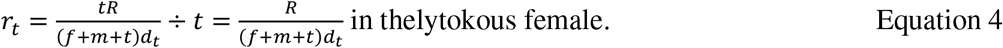

The condition under which the rate of increase in sexuals exceeds that of thelytoky is

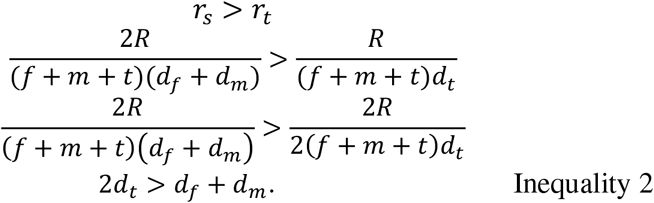

It is satisfied when *d*_*t*_ > *d*_*m*_ > 0 since *d*_*f*_ = *d*_*t*_. This is the same result as in the case of interlarval competition (inequality 1). Since the two independent models drew the same conclusion, it is very general and robust.

Model 2 corresponds to the situation where the male secures the same amount of resources as the female but does not consume them all himself, and the surplus is offered to the female and her offspring. Male behaviours such as territorial defence, monogamy, paternal care and nuptial gifts correspond to this situation. If the male acquires resources but does not contribute to the female’s reproduction, anisogamy loses to thelytoky (see Model 3 in the Supplementary Information).

### b) The “harmful male” hypothesis

In animals such as butterflies under weak male-male physical competition for mates, adult body mass is usually smaller in males than in females^26^ because males do not have heavy ovaries. In such species, the cost of growth would be lower for males. However, in species such as beetles under strong male-male physical competition^27^, male size and growth costs are often greater than those for females. The light eater male hypothesis cannot apply to such species, but males, because of their large size, can contribute in other ways.

#### Sexual conflict produces harmful males

In panmictic populations, an investment sex ratio of 1:1 is evolutionarily stable^4,11^. Even in this equilibrium, the number of spermatozoa is far greater than the number of eggs throughout the population. As a result, males can mate with many more females, even if all females are inseminated. Males force females to mate with them and disturb females to mate with other males. This sexual conflict^28^ leads to the evolution of harmful male traits (e.g., a spiny penis^29^, grasping organs^30^ and suckers^31^ that restrain females, and toxic seminal fluid^32^). These traits weaken females, and the weakened females increase their current reproductive investment and egg production. This allows harmful males to increase the number of eggs fertilised by their own sperm.

#### Males can immediately punish thelytoky invaders

Even under mutualism, derived asexuals (thelytoky), which abandon sex to escape from the male production cost, will invade sexual populations as cheaters. Thelytokous females threaten the niche of anisogamous females because once they emerge, they become a separate species that is reproductively isolated from the sexual population. Since newly evolved asexual females should be very similar to sexual females in phenotypes, males are unable to distinguish between the two^33^. Thus, the redundant male mating capacity is directed towards asexual females. Forced mating by harmful males reduces the fitness of asexual females^19,34,35^. When males are larger (more resource demand) than females (*d*_*f*_ = *d*_*t*_ < *d*_*m*_) and if harmful mating by males relatively reduces the survival of thelytokous females to *S*_*ah*_ (survival after harm, *S*_*ah*_ < 1), Models 1 and 2 are modified as follows:

Model 1

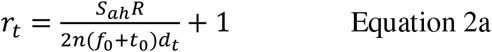

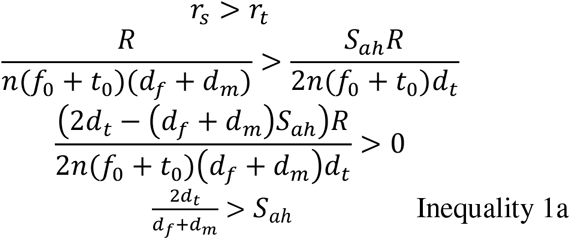

Model 2

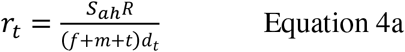

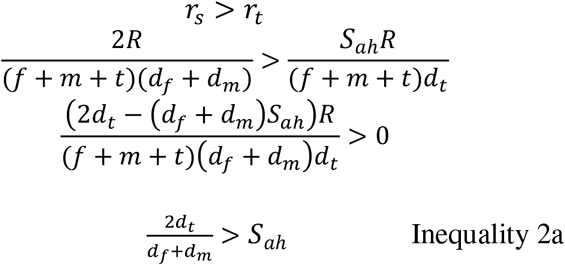

Again, the same conclusion is drawn from Model 1 (offspring competition) and Model 2 (adult competition). Inequalities 1a, 2a and Simulation 3 (Fig. 3) show that when male resource requirements (body size) are twice those of females (*d*_*f*_ = *d*_*t*_ = 1 and *d*_*m*_ = 2), harmful mating can protect anisogamy from thelytoky invasion if it reduces female survival by less than two-thirds (*S*_*ah*_ < 0.667, Fig. 3a). When there is no size difference (*d*_*f*_ = *d*_*t*_ = *d*_*m*_ = 1), the slight male harm (1% reduction in survival in thelytoky) is sufficient (Fig. 3b). Thus, the twofold benefit of anisogamy against the speculative twofold cost is unnecessary.

**Fig. 3.**
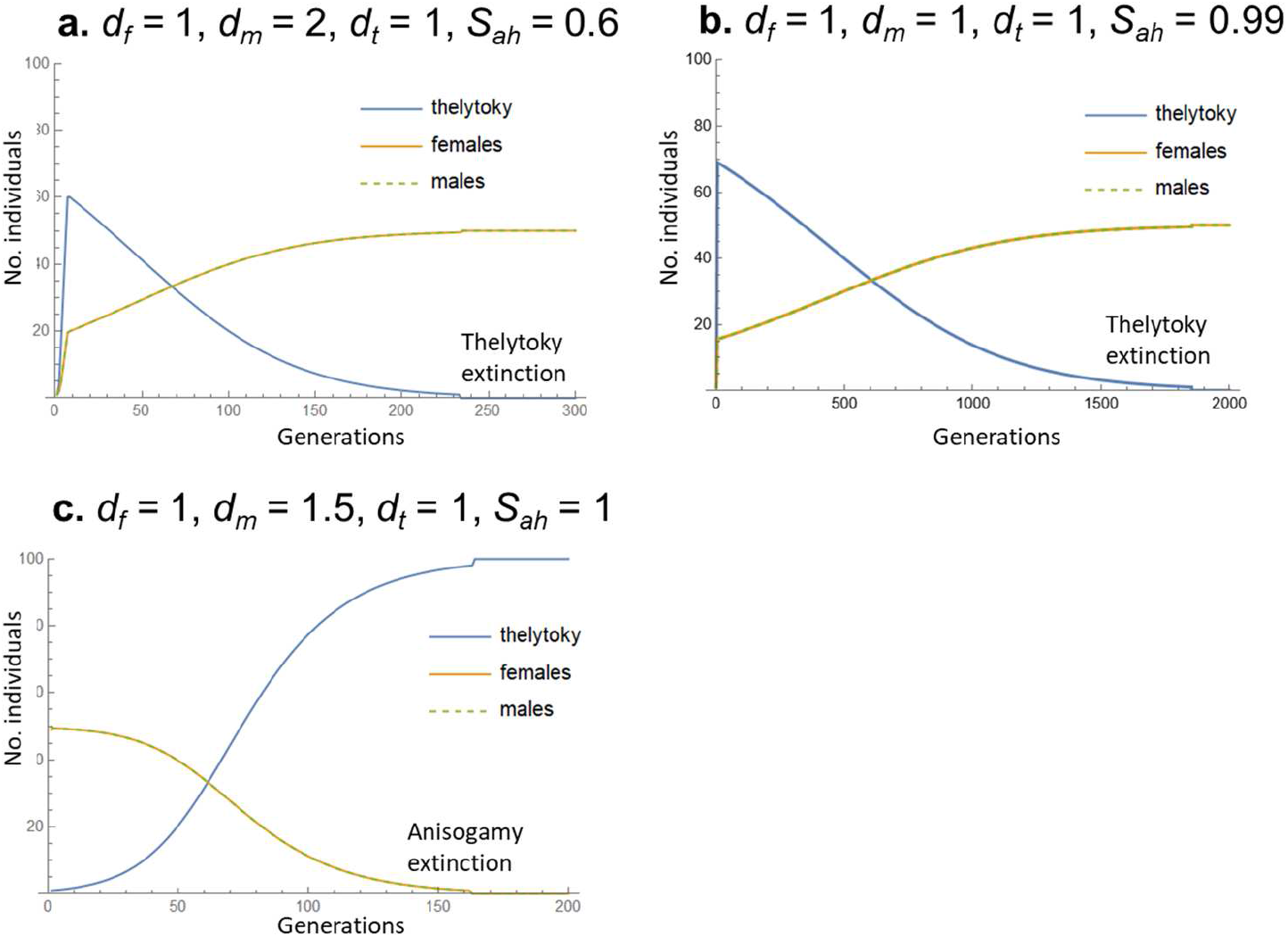
Results of simulation 3. When a mutant female abandoned sex and evolved into a new thelytokous species (female betrayal) in an anisogamous population with finite resources (carrying capacity = 100), the harmful males punished the invader and protected the anisogamous population. The thelytokous genotype had twice the reproductive rate of the sexual genotypes only at low density. *d*_*f*_, *d*_*m*_ and *d*_*t*_: growth costs (body size) in anisogamous females, males and thelytokous females, respectively. *S*_*ah*_: survival of thelytokous females after harmful mating. **a**. At a male/female size ratio of 2, 40% mortality due to harmful mating prevented thelytoky invasion. **b**. Without size dimorphism, only the 1% mortality was sufficient. **c**. When a mutant thelytokous female emerged, if males did not attack her (male betrayal), the sexual population was wiped out by the rapidly increasing thelytokous females. Thus, to block thelytoky invasion, both sexes must cooperate; females producing sons as bodyguards and males guarding the sexual population.

## Discussion

### Male production cost is the investment to maintain female-male mutualism

Mutualism is a mutually beneficial but mutually costly interaction between selfish players^22,36^. In this interaction, costs are transformed into necessary investments to realise cooperation. In anisogamy, females invest in large eggs and guarantee the presence of males by making half of their offspring male, and males invest in the mass production of sperm to ensure the fertilisation of eggs. Males make various contributions to maintain anisogamy, such as (1) reallocation of their unused resources to females or offspring and (2) sterile mating with thelytokous females to protect the sexual population.

Maintaining mutualism requires not only rewarding cooperators but also punishing betrayers^22,36^. Under sexual conflict^28^, there is a potential selection pressure for females to stop producing sons and evolve into asexual species and for males to cease mating with asexual females and choose only sexual females. However, if one betrays the other, the betrayer is punished; asexual mutant females are attacked by harmful males, and mutant males that abandon interspecific mating lead to the extinction of the sexual population to which they belong (Fig. 3). Because this mutual punishment system is not perfect, thelytoky sometimes evolved. However, they lose the long-term benefits of sex (see Fig. 1 of ^10^) and do not last long. In fact, thelytoky species are at the tip of phylogenetic trees that include sexual species, and their branches are short^37,38^. Although both sexes invest in maintaining anisogamy, researchers have focused only on female investment and traditionally called it “the twofold cost of male production”.

### “Nonproductive” males are essential to sustain anisogamy

In Models 1 and 2 and Simulation 2, the resource-saving anisogamy overcame thelytoky with a twofold increasing potential. Why does this occur? In our finite resource settings, not only the number of offspring born but also their resource demand for individual development determine the increasing rates (Equations 1∼4). Because they are equally competitive, thelytokous individuals cannot deprive resources from sexual individuals, meaning that resource saving becomes more important. In the thelytoky, the resources secured by *n* individuals can support a maximum of *n* individuals. If the survival rate during development is 80%, 0.8*n* individuals will reach adulthood. In anisogamy, if a 10% saving of resources by males (5% average between males and females) can increase the survival rate of anisogamy by 5%, 0.84*n* offspring will mature. Thus, only anisogamy can increase in frequency after saturation by replacing thelytoky (Fig. 2a). Although Model 2 assumed special male behaviour, such as paternal care or territorial defence, this is not always necessary. As shown in Model 1, it would work if the resources not used by the less demanding males are retrieved by females and increase their survival. This hypothesis is therefore applicable to a wide range of animals.

Anisogamous females pay the egg production cost, but thelytokous females also pay the same cost because the number of eggs laid by both species is the same. The only difference is that in anisogamy, half of the eggs are male. It is believed to incur a twofold male-production cost. However, after the population reaches carrying capacity, this does not impose such a high cost on the anisogamous population because thelytoky no longer express a twofold increasing rate. Consequently, anisogamy wins thelytoky if the males contribute even slightly to female reproduction. Therefore, contrary to the traditional view^2,3,6,7^, male nonproductivity is not a cost but a benefit for females, as it reduces resource use by not producing eggs. A comparative study^39^ shows that asexually reproducing species tend to thrive in harsh, low-density environments around their distribution range, whereas sexually reproducing species are more common in high-density environments, such as tropical rainforests containing highly coevolved communities. This means that thelytoky can only outperform anisogamy in populations well below carrying capacity, supporting our view that resource conservation through anisogamy is more important than high productivity in saturated populations.

### Asexual females are not asexual type of the same species

Traditional arguments^18,20,40^ regarding the cost of sex have considered thelytokous females derived from sexual populations as asexual morphs of the same species (belonging to a gene pool). However, once an asexual female arises in the sexual gene pool, the asexual female and the sexual individuals no longer share that gene pool (i.e., no gene flow from thelytokous to the sexual population), and they become separate species. Thus, the relationship between sexual and asexual individuals must be considered interspecific competition.

This is the actual reason why once the thelytokous mutant has appeared in a sexual population, the sexual genotype cannot suppress the increase in thelytoky. Anisogamy was able to beat ancestral isogamy despite the twofold cost of male production (Fig. 1) because they interact within the same sexual gene pool (isogametes also accept sperm). For example, even if some mothers produce offspring with a female-biased sex ratio to reduce the cost of males or to return to isogamy, the increased frequency of the female-producing allele X favours the male-producing allele Y because it increases the chance of male mating. This negative frequency-dependent selection suppresses the increase in X. However, the gene T that makes thelytokous females is in its own gene pool (asexual population) and is not subject to dynamics within the sexual gene pool, such as frequency dependence or kin selection. Kawatsu^19^ set up asexual females to mate with males and produce sexual females and males (reproductive isolation not yet established) to pull asexual females back into the sexual gene pool. There, negative frequency dependence (i.e., an increase in females favouring males) could work, and thus, Kawatsu’s^19^ conclusion that harmful males prevent the invasion of thelytoky would be valid. However, although there are some examples in insects^41-44^, facultative parthenogenesis is extremely rare and difficult to generalise.

Therefore, the only way to stop thelytoky invading anisogamous populations is either (1) thelytoky self-destructing within their own gene pool, or (2) males attacking asexual females from outside thelytoky gene pool. The “light-eater male” and “harmful male” hypotheses correspond to (1) and (2) respectively.

### Is harmful mating evolvable?

Rankin^20^ concluded that harmful males that mate with thelytokous females cannot evolve because such infertile mating reduces the male’s own fitness, and thus, males choose sexual females as mates. However, his argument is based on the miscalculation of relatedness^45^ between males and thelytokous females (see the Supplementary Information). We consider that harmful mating is evolvable even if it does not produce viable offspring. It can be selected in two ways: 1) Direct fitness increases in mating males; if the males have already left offspring in the sexual population, attacking thelytoky increases their individual fitness by the same mechanism as the evolution of paternal care of offspring^22^. 2) Indirect fitness increases via multilevel or kin selection^46,47^; even if the males have not yet reproduced, harmful mating with asexual females helps their kin in the sexual gene pool and increases their inclusive fitness by the same mechanism as the evolution of sterile soldiers in eusocial insects^22,47^. Harmful mating is observed in some asexual fish, where diploid self-developing eggs are fertilised by sperm, and they become triploid, resulting in abnormal embryonic development, cancer, infertility and other adverse effects, often leading to death^48-50^.

### Preadaptations ease the evolution of harmful males

Males have often acquired weapon traits (e.g., large body size and horns)^21^ through intrasexual selection and harmful traits through sexual conflict^28^. These male traits become preadaptations to mate with asexual females, so additional trait evolution and its extra costs are unnecessary. Each time a thelytoky mutant appears, many harmful males immediately punish the invaders.

### Counteradaptations occur only in sexual females

Under sexual conflict, antagonistic coevolution known as “chase away”^51^ endlessly occurs between male harmfulness and female resistance so that male harmfulness rapidly evolves into another type of harmfulness. Sexual females can counterevolve by combining resistance genes that occur in different individuals in the gene pool (i.e., the Red Queen process^13,14^), but asexual females, which can only access mutations that occur in their own lineage, cannot follow the rapid evolution of males. In such cases, harmful males will reduce only the fitness of asexual females without harming the sexual population. Some empirical studies in fish and reptiles have shown that the presence of males negatively affects not conspecific females but thelytokous females of related species^34,35^. Therefore, harmful males may be still mutualistic with conspecific sexual females.

## Conclusion

This study provides a paradigm shift in the arguments about the twofold cost of male production. Anisogamy is not male parasitism on females but mutualism between the sexes. Previous authors have looked for short-term benefits of sex that outweigh the twofold costs but have not found them. Instead, we looked for mechanisms that prevent thelytoky from expressing their twofold reproductive potential. Light eater males and harmful males are just two examples, and any other mechanism by which males reduce the thelytoky’s performance could maintain anisogamy.

Anisogamy flourishes because, in many cases of finite resources, male production costs are less than twofold as high. Promising future studies to test our hypotheses include 1) whether males consume fewer resources than females and whether redundant resources are reused by sexual populations, 2) whether harmful males reduce the fitness of asexual females more than sexual females, and finally 3) whether the evolution of thelytoky tends to be suppressed in lineages in which males develop weapons^21^ or sexual conflict traits^28^.

## Methods

### Computer simulations of population dynamics

A program was written using Mathematica for Windows ver. 13.0 (Wolfram language).

#### (1) Invasion of anisogamy into an isogamous population (Fig. 1)

Considering that a hypothetical mutant gene Y that produces small gametes (i.e., sperm) arises, Y was dominant over the wild-type gene X, which produces isogametes. Individuals with genotype XY become males and produce X sperm and Y sperm in a 1:1 ratio. Wild-type individual XX produces isogamete X, which maintains isogamy if it is fertilised by another isogamete X of a different mating type^6,25^. Isogamete X fertilised by X and Y sperm become anisogamous females and males, respectively. Sperm could be fused to any isogamete in the population since it was a new mating type, i.e., the population sex ratio is extremely isogamous female biased. The anisogamous zygotes were smaller than the isogamous zygotes due to the smaller sperm resources but were able to survive and develop because an absolute requirement for the evolution of anisogamy is that the embryo did not die when one parent reduced the size of the gametes. This condition is realised by the theory of Parker *et al*.^5^ that the variation in gamete size gradually expanded by disruptive selection or by the “inflated isogamy” hypothesis of Yasui and Hasegawa^10^ that an increase in the size of the isogametes would precede the evolution of sperm. There are no YY individuals because sperm-to-sperm fusion is not viable. In generation 1, an isogamous population (carrying capacity = 100) consisted of 99 wild-type isogamous individuals (genotype XX) and one male (XY). Mating between two isogamous parents will produce 4 isogamous daughters (XX), while mating between an isogamous female and a male will produce 2 isogamous daughters (XX) and 2 sons (XY). The isogamete fertilised by the sperm is now called an egg. Sperm have priority in fertilising eggs because they move faster and are more numerous than isogametes; one male fertilises all eggs of *dp* females. For example, if *dp* = 2, the first male fertilises 8 eggs from 2 females, leaving 4 daughters and 4 sons. The remaining 97 females make 48 isogamous pairs (one unmated due to monogamy), and their isogametes make 192 females, but with a probability of 1 *k*, resulting in unfertilisation. If the male mating capacity (number of males × *dp*) exceeds the number of females, the excess males cannot mate. The two phenotypes (isogamous female and male) are equally competitive with respect to the carrying capacity of the population. The simulation was continued until equilibrium was reached.

#### (2) The light eater male hypothesis (Fig. 2)

Equal competition between juveniles under finite resources determines the amount of available resources for each individual. Males consume fewer resources than females, and the surplus is given to sexual females. At low densities, the potential increasing rates of anisogamy (*r*_*s*_: Equation 1) and thelytoky (*r*_*t*_: Equation 2) calculated in Model 1 are larger than 2 and 4, respectively. In such cases, we set an anisogamous pair to produce 2 females and 2 males per generation, while a thelytokous female produces 4 females per generation. If the population reached carrying capacity, the three phenotypes (sexual female, thelytokous female and male) compete according to their own increasing potential (*r*_*s*_ or *r*_*t*_). Thus, anisogamy entails the twofold cost of male production only at low densities.

#### (3) The harmful male hypothesis (Fig. 3)

The parameter *S*_*ah*_, the survivorship of thelytokous females after mating with harmful males, was added to the simulation program (2).

## Acknowledgements

Y. Y. thanks Kazuya Akimitsu for his encouragement on submissions. Tatsumi Kudo and Kazuya Kobayashi provided literature information. **Funding:** This work was partly supported by Grants-in-Aid from the Ministry of Education, Culture, Sports Science and Technology of Japan to Y. Y. (nos. 26440241, 19K06839 and 21K19116) and to E. H. (nos. 18H02502 and 19H0296400).

## Author contributions

YY and EH conceived the original idea and contributed to the logical construction through thorough discussions. YY made the models and performed the simulations. YY mainly wrote the manuscript, and EH checked it for logics and presentations. As a result, both authors contributed equally to this article.

## Competing interests

The authors declare no potential conflicts of interest. We followed all ethics guidelines of the journal.

## Code availability

Codes for the simulations are provided as supplementary information.

## Supplementary information

### Model 3: Adult males secure resources but do not provide them to offspring

The numbers of adult individuals of anisogamous females, males and thelytokous females are *f, m* and *t*, respectively. Adults are equally competitive, so the resource is divided equally between the three parties into 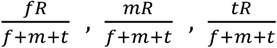. Sexual females reproduce using only their own resources, 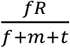. This is because there is no paternal investment. One juvenile sexual female, one male and one thelytokous female need resources *d*_*f*_, *d*_*m*_ and *d*_*t*_, respectively, to grow to adulthood. The number of offspring reaching adulthood in a sexual population is 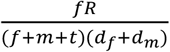 pairs at a sex ratio of 1:1. The number of offspring reaching adulthood in the thelytokous population is 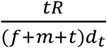. The rate of increase per generation is

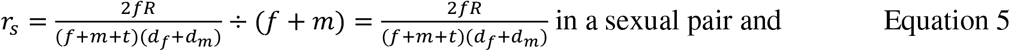

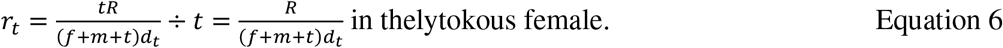

The condition under which the rate of increase in sexuals exceeds that of thelytoky is

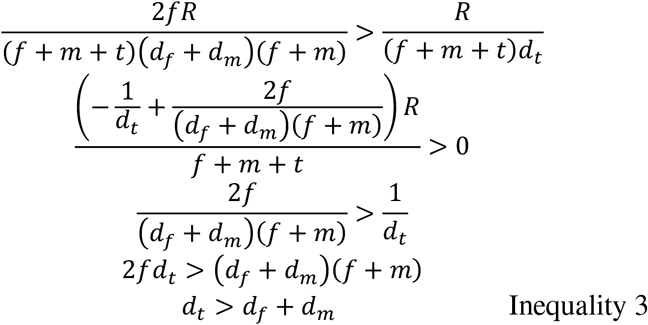

because of the 1:1 sex ratio *f* = *m*. This means that the maintenance of anisogamy requires the condition where one thelytokous female needs more resources than the sum of the needs of one sexual pair. This is unrealistic.

### One misconception about harmful males

Rankin^20^ has argued that males harmful to thelytokous females cannot evolve because mating with asexual females does not increase male fitness, whereas males that choose sexual females are favoured. This author also suggested that sexual females could instead punish thelytokous females, but several key assumptions in that paper were inappropriate. For example, it assumed that the relatedness of sexual males to asexual females is r = -1. In studies of eusocial insects, the regression relatedness^45^ of a donor (e.g., worker) to a recipient (e.g., queen) is calculated as the slope of the regression of the recipient’s degree of having the focal allele on the donor’s degree (see Fig. S1a for details). Importantly, when calculating relatedness, the donor and recipient must be in the same gene pool because the average frequency of the focal allele in the population is required to control the probability that the focal allele is shared by chance. Therefore, the calculation^18,20^ across different gene pools (or two species) is wrong because the average frequencies are different. Moreover, even if the two species are forcibly set as a single gene pool, the relatedness is not -1. If we focus on several independent loci, two related individuals tend to have a particular allele to the same degree (none, heterozygote or homozygote) on each locus, and then the regression line should have a positive slope (Fig. S1a). However, all the thelytokous females do not have focal alleles or have them to the same degree as their clonal common ancestor. Consequently, the regression line has no slope (Fig. S1b), and the relatedness becomes zero. Some authors^18,20^ have argued whether male punishment of asexual females can evolve as a spiteful behaviour that requires a negative relatedness between players, but their arguments are based on such miscalculations and are thus invalid. Furthermore, another suggestion in that paper^20^ (harmful females to thelytokous females) is unlikely to evolve because if sexual females attack thelytokous females, sexual females would pay considerable costs (e.g., being wounded or killed). Thelytokous females that arose from sexual females would have the same physiological mechanisms, fighting abilities and niche as the sexual ancestor in addition to the twofold increasing rate. Thus, sexual females could not win this competitor. In our view, harmful males are able to evolve not as spite but by their own fitness contribution to the sexual population (see text).

**Fig. S1.**
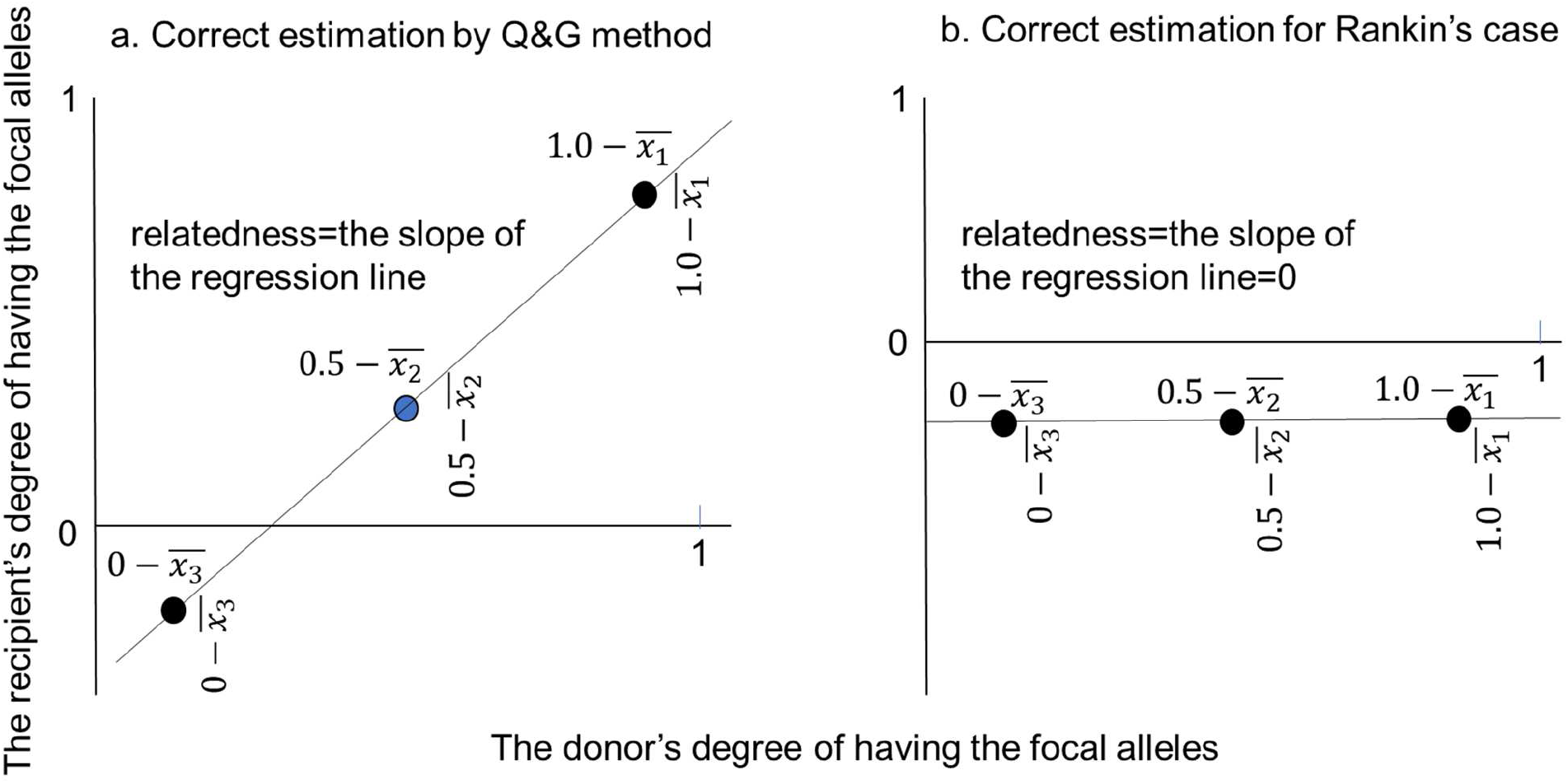
Regression relatedness between two individuals. a) An example of the calculation of the regression relatedness by the Queller and Goodnight^45^ method. In a single gene pool, there are multiple (3 in this example; *x*_1_, *x*_2_ and *x*_3_) loci shared by a donor and a recipient of some social behaviour (e.g., altruism). The donor (2n) has a focal allele on each locus at three frequencies: 0 (none), 0.5 (heterozygote) or 1.0 (homozygote). The recipient also has its own frequencies. If both individuals are related, this value will tend to coincide for each locus. Then, the recipient’s frequencies are plotted on the donor’s frequencies after subtracting the population average frequency of the focal allele (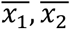 and 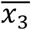) to correct allele sharing by chance. The slope of the regression line is an estimate of the relatedness of the donor to the recipient. b) Rankin^20^ set the relatedness of a sexual male (donor) to an asexual female (recipient) as -1.0, but this is incorrect (see text).

## Notes

### Competing Interest Statement

The authors have declared no competing interest.

